# ADP-ribosylation drives the repair of topoisomerase IIIβ-induced nucleic acid damage

**DOI:** 10.64898/2026.11.19.767422

**Authors:** Sourav Saha, Tessa Maidens, Victor Cai, Yijun Su, Jiji Chen, Nina C. Lee, Tapan K. Maity, Lisa M. Jenkins, Linying Zhu, Bo Wu, Weixing Zhao, Gerald M. Wilson, Yves Pommier, Yilun Sun

## Abstract

Abortive DNA and RNA-linked covalent cleavage complexes formed by the topoisomerase IIIβ can disrupt nucleic acid metabolism. Here, we describe poly(ADP-ribosyl)ation (PARylation) sites on TOP3B cleavage complexes (TOP3Bccs) and demonstrate their relevance for the repair of both DNA- and RNA-linked TOP3Bccs. We show that persistent TOP3Bccs are rapidly PARylated by PARP1 and de-PARylated by poly(ADP-ribose) glycohydrolase (PARG). PARylation recruits Flap Endonuclease 1 (FEN1), which excises TOP3Bccs. We also show that PARG inhibition recruits the deubiquitylating enzyme (DUB) USP10 to promote TOP3Bcc deubiquitylation and prevent their proteasomal degradation. Together, this study identifies a dynamic PARylation-mediated regulatory pathway coordinating the repair of abortive TOP3Bccs on DNA and RNA. This pathway defines a critical mechanism of TOP3Bcc homeostasis and may influence cancer cell responses to TOP3B-targeting agents.

## INTRODUCTION

Human topoisomerases comprise a highly specialized family of six enzymes (TOP1, TOP1MT, TOP2A, TOP2B, TOP3A and TOP3B) to resolve topological problems in DNA/RNA such as supercoils, knots and catenanes that arise during replication, transcription, translation and recombination (1–3). Topoisomerases act by cleaving the DNA or RNA backbone and forming protein-DNA/RNA covalent catalytic intermediates (at the 3’-end for TOP1 and TOP1MT and 5’-end for TOP2 and TOP3) termed topoisomerase cleavage complexes (TOPccs) (1–3). Under physiological conditions, TOPccs are short-lived, readily reversible and undetectable. However, endogenous DNA lesions, including oxidized bases, nicks, abasic sites, and misincorporated ribonucleotides, as well as anticancer topoisomerase inhibitors such as topotecan and irinotecan for TOP1 and etoposide, doxorubicin, and amsacrine for TOP2 prevent religation and trap otherwise transient TOPccs. The resulting protein-linked nucleic acid breaks interfere with essential DNA and RNA transactions and are highly cytotoxic (4–9).

Human topoisomerase IIIβ (TOP3B) is unique among topoisomerases because it acts on both DNA and RNA (10–15). TOP3B relaxes hyper-negatively supercoiled DNA and resolves intertwined DNA and RNA structures (knots and catenanes) by transiently cleaving a single nucleic acid strand, forming a covalent 5′-phosphotyrosyl TOP3B cleavage complex (TOP3Bcc) permitting passage of another strand before resealing the break (10–14). Although TOP3Bccs are normally transient, they can be aberrantly stabilized under pathological or experimentally induced conditions including: 1) loss/depletion of TDRD3 (16), the scaffolding protein for TOP3B; 2) expression of the autism-link*ed de n*ovo mutant variant C666R TOP3B (17) or religation-defective “self-trapping” mutant R338W TOP3B (12); 3) treatment with the TOP1 inhibitor camptothecin and the splicing inhibitor pladienolide-B (18); and 4) treatment with recently discovered TOP3B trapping agents thiacyanine (NSC96932) and symmetrical bisacridine (NSC690634) (19). Persistent accumulation of TOP3Bccs on both DNA and RNA induce R-loop (12,16,18,19), ribosome collision (17) and marked cytotoxicity in rapidly proliferating cancer cells (12,16,18,19).

The mechanisms that repair TOP3Bccs remain largely understudied. We previously reported that TOP3Bccs can be repaired through the ubiquitin-proteasome pathway and tyrosyl-DNA phosphodiesterase 2 (TDP2) (12). TRIM41, a ubiquitin E3 ligase, ubiquitylates and primes TOP3Bccs for proteasomal degradation (12). These findings established that cells possess dedicated mechanisms for degrading abortive TOP3Bccs, but whether additional post-translational modifications regulate the recognition, processing, or stability of DNA- and RNA-linked TOP3Bccs remains unknown.

Poly(ADP-ribosyl)ation (PARylation) is a rapid and reversible post-translational modification playing central roles in the cellular response to nucleic acid damage. Poly(ADP-ribose) polymerases (PARPs), particularly PARP1 use NAD⁺ to synthesize poly(ADP-ribose) chains on themselves and other proteins, thereby altering protein activity and promoting the recruitment of repair factors to sites of damage. PARylation is highly dynamic and rapidly reversed by poly(ADP-ribose) glycohydrolase (PARG) (20–25). Here, we identify PARP1-mediated PARylation as a key regulator of TOP3Bcc homeostasis. We show that both DNA- and RNA-linked TOP3Bccs are PARylated and that dynamic PARylation/dePARylation promotes their resolution by recruiting flap endonuclease 1 (FEN1). In contrast, persistent PARylation caused by PARG inhibition recruits the deubiquitylase USP10, and stabilizes TOP3Bccs. Together, these findings establish PARylation as a dynamic molecular switch that coordinates the repair of TOP3B-associated nucleic acid damage.

## RESULTS

### 1. Dynamic PARylation regulates cellular TOP3Bcc homeostasis

To discover novel cellular pathways involved in TOP3Bcc repair, we performed a high-throughput chemical screen in cultured human cancer cells against the TOP3B poison NSC690634(19). PARP inhibitors including olaparib, rucaparib and niraparib enhanced the cellular response to TOP3B trapping (**Figure 1A**). This result is consistent with our prior observation that aberrant TOP3Bccs caused by loss of the TOP3B scaffolding protein TDRD3 activates PARP1 and enhances the interaction between TOP3B and PARP1 in an earlier study (16). Together with our chemical screening results, these observations prompted us to determine whether PARylation directly regulates TOP3Bcc formation and resolution.

**Figure 1.**
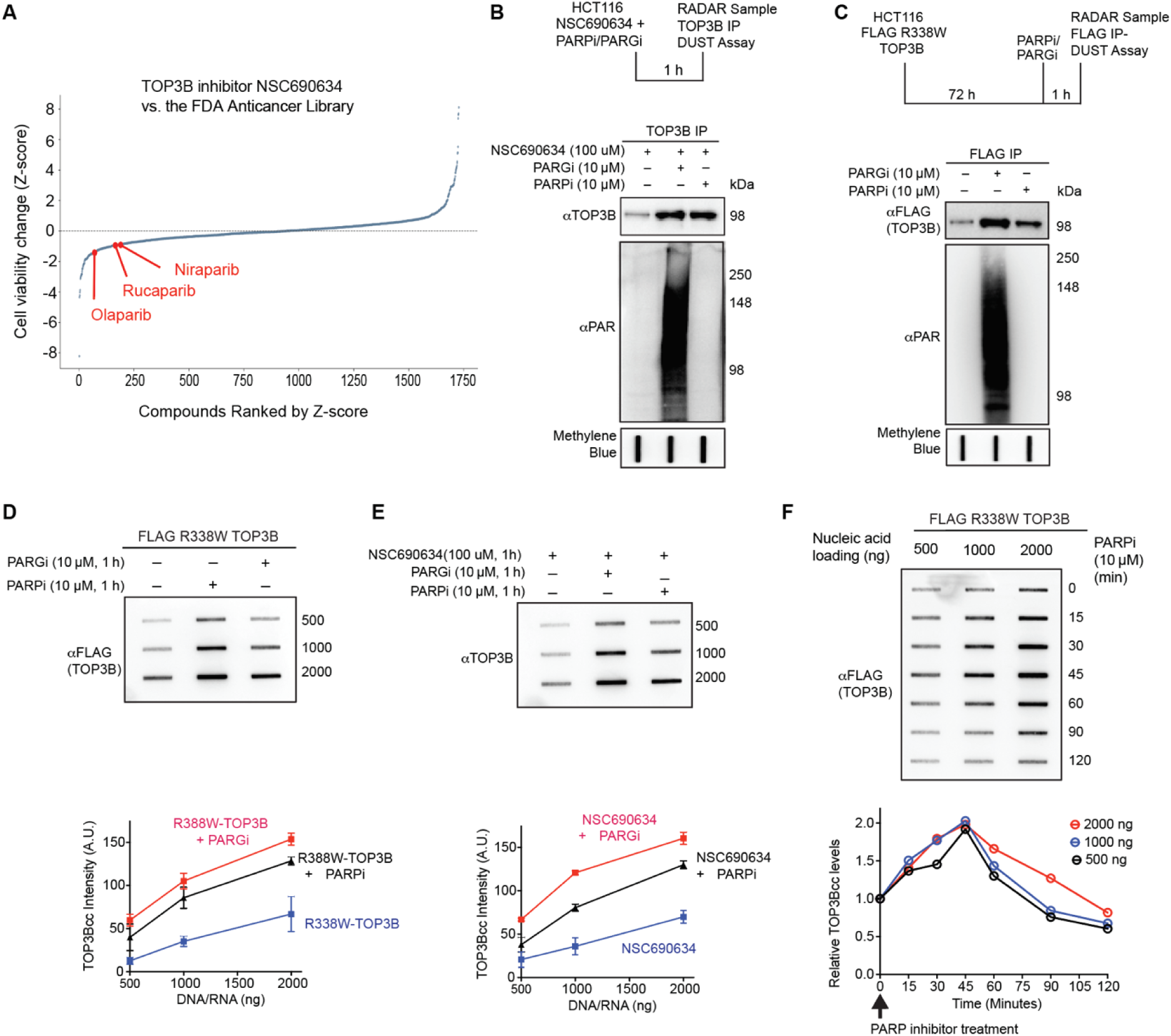
Dynamic PARylation regulates cellular TOP3B cleavage complex homeostasis. **A.** High-throughput drug combination screening identifies PARP inhibitors as candidate sensitizers to TOP3B inhibition. DMS114 small cell lung cancer (SCLC) cells were screened against an anticancer compound library in the absence or presence of the TOP3B inhibitor NSC690634. Cell viability was measured using CellTiter-Glo, and differences in viability between single-agent and combination treatments were standardized as Z-scores. Each dot represents an individual compound ranked by its Z-score. More negative Z-scores indicate greater apparent sensitization to NSC690634. **B.** Endogenous TOP3Bccs are PARylated. HCT116 cells treated with the TOP3B inhibitor NSC690634 (100 μM, 1 h) were additionally treated with PARPi or PARGi. Endogenous TOP3Bccs isolated by RADAR were immunoprecipitated with anti-TOP3B antibody and analyzed for PARylation by immunoblotting. **C.** Cellular TOP3Bccs are PARylated. HCT116 cells expressing FLAG-tagged R338W TOP3B were treated with PARP inhibitor (PARPi; 10 μM, 1 h) or PARG inhibitor (PARGi; 10 μM, 1 h), as indicated. TOP3Bccs isolated by the RADAR assay were immunoprecipitated with anti-FLAG antibody, digested with micrococcal nuclease, and analyzed by immunoblotting for PAR and FLAG-TOP3B. Methylene blue staining shows nucleic-acid loading. **D.** PARylation is required for TOP3Bcc resolution, whereas persistent PARylation following PARG inhibition results in TOP3Bcc accumulation. HCT116 cells expressing FLAG-tagged R338W TOP3B were treated with PARPi or PARGi as indicated, and TOP3Bccs were quantified by RADAR slot blotting with anti-FLAG antibody. Unless otherwise indicated, quantification data from three independent experiments are presented as mean ± SD. A.U., arbitrary units. **E.** Dynamic PARylation is required for efficient endogenous TOP3Bcc resolution. TOP3Bccs induced by NSC690634 were quantified following PARPi or PARGi treatment by RADAR slot blotting with anti-TOP3B antibody. **F.** PARP inhibition causes rapid accumulation of TOP3Bccs. HCT116 cells expressing FLAG-TOP3B R338W were treated with PARPi for the indicated times, and TOP3Bcc levels were measured by RADAR slot blotting.

We next asked whether endogenous TOP3Bccs generated by pharmacological TOP3B trapping are PARylated by performing modified RADAR (rapid approach to DNA adduct recovery) assays (12,16,26) in HCT116 cells treated with NSC690634 with or without PARP or PARG inhibitors. PARylation of endogenous TOP3Bccs was most readily detected following PARG inhibition with PDD00017273 (27) and was suppressed by PARP inhibition (**Figure 1B**). These findings demonstrate that drug-induced endogenous TOP3Bccs are dynamically PARylated.

To further establish that cellular TOP3Bccs are PARylated, we performed RADAR assays in HCT116 cells expressing the self-trapping FLAG-tagged R338W TOP3B (12), immunoprecipitated (IP) with anti-FLAG antibody, digested with micrococcal nuclease and immunoblotted with anti-PAR antibody. PARylated TOP3Bccs were readily detectable following inhibition of PARG (**Figure 1C**). Conversely, treatment with the PARP inhibitor (PARPi) olaparib (28,29) abolished the TOP3Bcc PARylation signal. These results demonstrate that TOP3Bccs undergo rapid and reversible PARylation in cells.

In parallel, we examined whether PARylation affects the cellular abundance of TOP3Bccs. Modified RADAR assays showed that PARP inhibition increased the accumulation of FLAG-TOP3B R338W cleavage complexes compared with untreated cells, whereas PARG inhibition produced a greater increase (**Figure 1C**). A similar pattern was observed for endogenous TOP3Bccs induced by NSC690634, with inhibition of either PARP or PARG increasing TOP3Bcc levels and PARG inhibition producing the more pronounced effect (**Figure 1B**).

To further quantify the role of PARylation in TOP3Bcc homeostasis, we performed slot blotting with RADAR samples prepared from HCT116 cells expressing R338W TOP3B with or without PARPi or PARGi treatment. Both PARP and PARG inhibition resulted in increased TOP3Bccs (**Figure 1D**). Likewise, slot blotting showed that both PARP and PARG inhibition increased endogenous TOP3Bcc accumulation following NSC690634 treatment, with PARG inhibition producing a particularly marked increase (**Figure 1E**). Thus, inhibition of PAR synthesis and inhibition of PAR turnover both perturb TOP3Bcc resolution, indicating that dynamic PARylation, rather than simply the presence or absence of PAR, is required for efficient TOP3Bcc homeostasis.

Because PARP inhibition produced only modest effects on TOP3Bccs compared to PARG inhibition, we examined the kinetics of TOP3Bcc accumulation following PARP inhibition in HCT116 cells expressing FLAG-TOP3B R338W. TOP3Bccs increased rapidly after PARP inhibition and reached a maximum at approximately 45 min (**Figure 1F**). These kinetics suggest that both PARP and PARG inhibitors enhance TOP3Bccs, and PARylation a reversible process.

### 2. PARylation regulates both the DNA and RNA TOP3Bccs

Because TOP3B is a dual-activity topoisomerase acting both on DNA and RNA (12–14), we tested whether PARylation affects TOP3Bccs on both nucleic acid substrates. To distinguish DNA and RNA-linked TOP3Bccs we used our established nuclease-based RADAR assay strategy (12,16) (**Figure 2A**). Isolated RADAR samples from HCT116 cells expressing FLAG-tagged R338W TOP3B were digested with DNase I or a mixture of RNase A and RNase T1 to isolate RNA- or DNA-linked TOP3Bccs, respectively. Equal amounts of the resulting RADAR samples were immunoprecipitated using FLAG antibody and analyzed by immunoblotting for PARylated DNA- and RNA-linked TOP3Bccs. Both the DNA- and RNA-linked TOP3Bccs were PARylated, with the PAR signal becoming readily detectable following PARG inhibition (**Figure 2B**). PARG inhibition also increased the abundance of TOP3Bccs on both DNA and RNA, indicating that persistent PARylation stabilizes TOP3Bccs irrespective of the nucleic acid substrate (**Figure 2B**).

**Figure 2.**
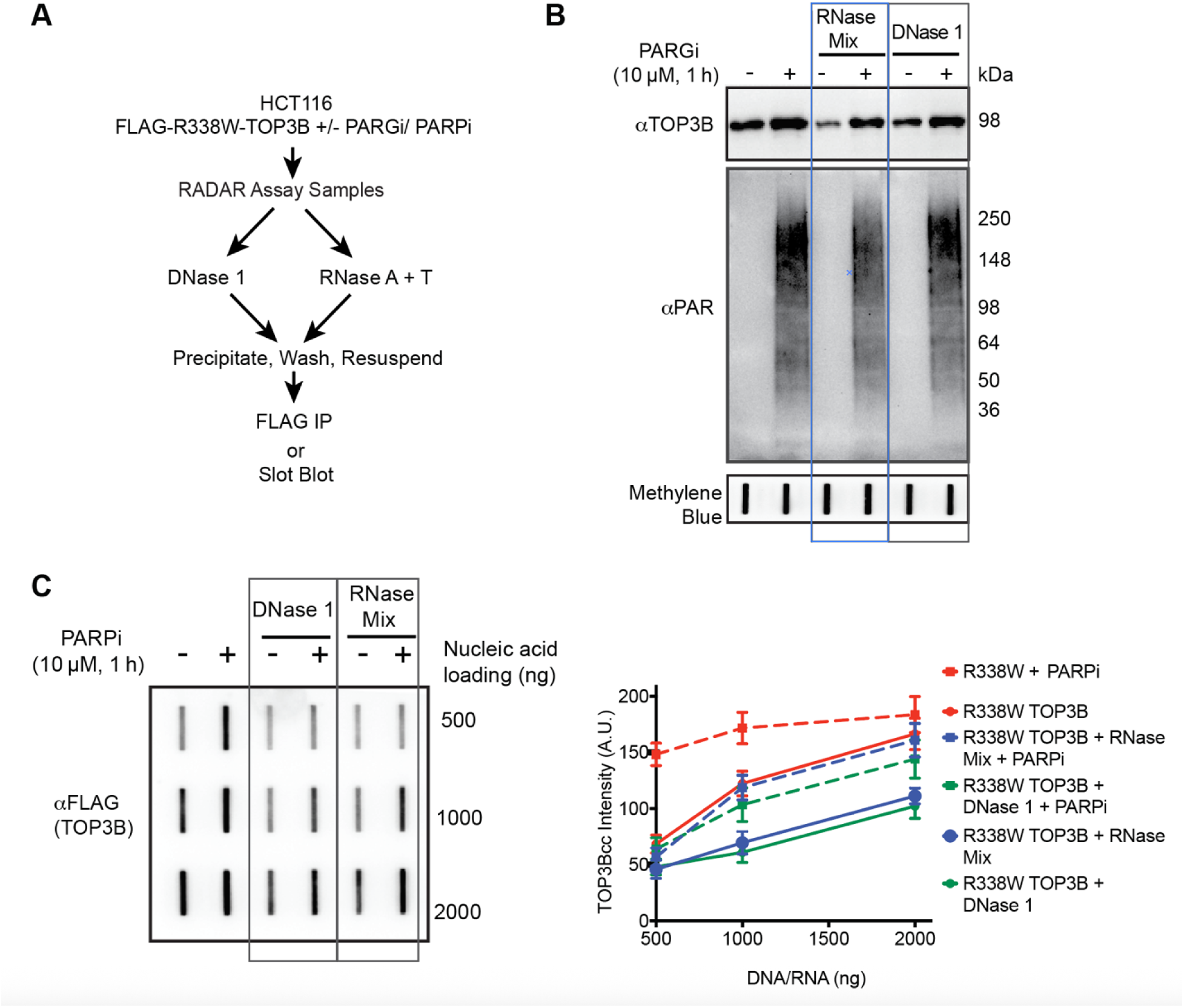
PARylation promotes the resolution of both DNA- and RNA-linked TOP3B cleavage complexes. **A.** Experimental strategy for distinguishing DNA- and RNA-linked TOP3Bccs. RADAR samples were digested with DNase I or a mixture of RNase A and RNase T1 to selectively analyze RNA- or DNA-linked TOP3Bccs, respectively. **B.** Both DNA- and RNA-linked TOP3Bccs are PARylated. HCT116 cells expressing FLAG R338W TOP3B were treated with PARGi (10 μM, 1 h). RADAR samples were treated with DNase 1 or RNase A/ RNase T1, immunoprecipitated with anti-FLAG antibody, and analyzed by immunoblotting for PAR and FLAG-TOP3B. **C.** PARP inhibition increases both DNA- and RNA-linked TOP3Bccs. HCT116 cells expressing FLAG-tagged R338W TOP3B were treated with PARPi (10 μM, 1 h), followed by selective nuclease digestion and RADAR slot blotting with anti-FLAG antibody. **D.** Quantification of the data shown in panel C from three independent experiments. Data are mean ± SD. A.U., arbitrary units.

We next examined the effect of PARP inhibition on DNA- and RNA-linked TOP3Bcc levels. DNA- and RNA-containing RADAR samples were prepared from HCT116 cells expressing FLAG-TOP3B R338W with or without PARP inhibitor treatment and analyzed by slot blotting. PARP inhibition increased TOP3Bcc accumulation in both the DNA and RNA fractions compared with untreated controls (**Figures 2C and 2D**). Altogether, these results demonstrate that both DNA- and RNA-linked TOP3Bccs undergo transient PARylation and that dynamic PARylation is required to maintain the homeostasis of TOP3Bccs on both nucleic acid substrates.

### 3. ADP-ribosylation of TOP3B at E438 and T488 determines TOP3Bcc homeostasis

To determine whether TOP3B can be directly ADP-ribosylated, recombinant TOP3B was incubated with recombinant PARP1, activated DNA and NAD⁺. Immunoblotting with an anti-PAR antibody confirmed that PARP1 ADP-ribosylates TOP3B *in vitro* (**Figure 3A**). Mapping of these ADP-ribosylation sites by LC-MS/MS (**Figure 3B**) identified nine ADP-ribosylated amino acid residues on recombinant TOP3B (**Figure 3C**).

**Figure 3.**
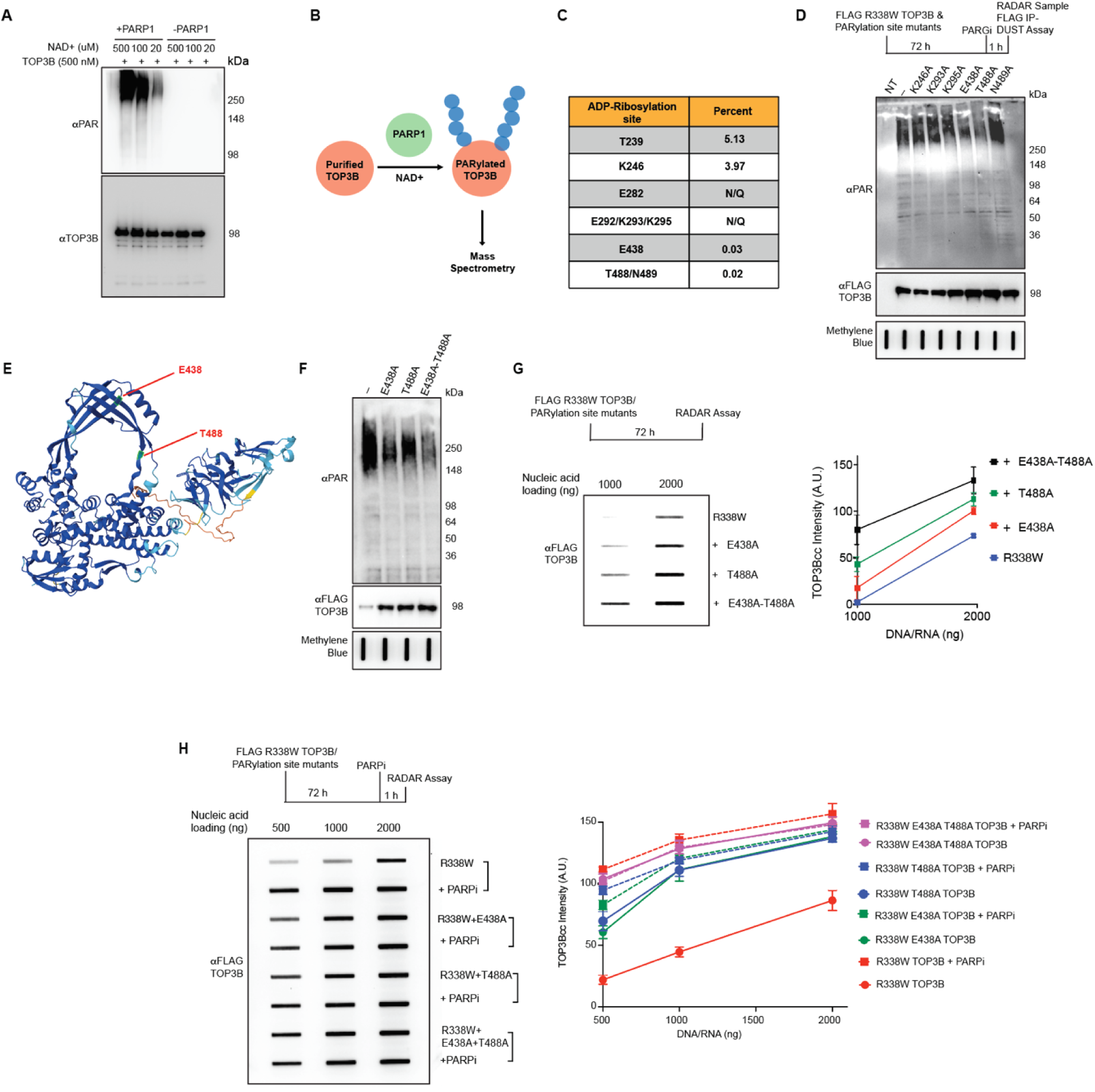
E438 and T488 are major TOP3B ADP-ribosylation sites required for TOP3Bcc homeostasis. **A.** PARP1 ADP-ribosylates TOP3B *in vitro*. Recombinant TOP3B was incubated with recombinant PARP1, activated DNA, and indicated concentrations of NAD⁺ for 20 min. Reactions were analyzed by immunoblotting with anti-PAR and anti-TOP3B antibodies. **B.** Workflow for proteomic mapping of ADP-ribosylated residues by LC-MS/MS following *in vitro* PARylation of recombinant TOP3B and digestion with trypsin/LysC or chymotrypsin. **C.** Identified TOP3B ADP-ribosylation sites and relative abundance. N/Q, no quantification data available. **D.** E438A and T488A mutations reduce cellular TOP3Bcc PARylation. HCT116 cells expressing FLAG R338W TOP3B or the indicated TOP3B mutants were treated with PARGi. TOP3Bccs isolated by RADAR were immunoprecipitated with anti-FLAG antibody and analyzed for PARylation by immunoblotting. **E.** AlphaFold predicted structure of human TOP3B (AlphaFold DB: AF-O95985-F1) highlighting glutamic acid residue 438 (E438) and threonine residue 488 (T488), ADP-ribosylation sites identified by LC-MS/MS. **F.** Combined mutation of E438 and T488 further reduces TOP3B PARylation. FLAG R338W TOP3B or TOP3B mutant proteins R338W/E438A, R338W/T488A, or R338W/E438A/T488A were analyzed following PARG inhibition as in panel D. **G.** Mutation of major TOP3B ADP-ribosylation sites promotes TOP3Bcc accumulation. TOP3Bcc levels were measured by RADAR slot blotting in cells expressing the indicated TOP3B mutants. **H.** PARP inhibition does not enhance the accumulation of cleavage complexes of ADP-ribosylation-deficient TOP3B mutants.

To determine which of these residues contribute most strongly to cellular TOP3B PARylation, we individually substituted candidate residues with alanine in the R338W TOP3B background and analyzed TOP3Bcc PARylation by modified RADAR assay following PARG inhibition. Among the mutants tested, mutations of glutamic acid E438 and threonine T488 showed the most pronounced reduction in PARylation (**Figure 3D**). **Figure 3E** represents the AlphaFold predicted structure of TOP3B (AlphaFold DB: AF-O95985-F1) (30,31) and shows that E438 and T488 are on the TOP3B protein surface. We therefore generated an E438A/T488A double mutant in the R338W background. The triple mutant (R338W/E438A/T488A) exhibited a greater PARylation defect than either double mutant (R338W/E438A or R338W/T488A) or the R338W single mutant (**Figure 3F**), indicating E438 and T488 as major ADP-ribosylation sites of TOP3B.

We next examined whether loss of ADP-ribosylation at these sites alters cellular TOP3Bccs. HCT116 cells expressing R338W TOP3B, R338W/E438A TOP3B, R338W/T488A TOP3B or R338W/E438A/T488A TOP3B were analyzed by RADAR slot blotting. All three PARylation-deficient mutants showed increased TOP3Bcc accumulation relative to R338W TOP3B, with the R338W/E438A/T488A triple mutant producing the highest TOP3Bcc levels (**Figure 3G**). These findings implicate ADP-ribosylation at E438 and T488 for TOP3Bcc resolution.

To further test whether these residues mediate the effect of PARP activity on TOP3Bcc homeostasis, we examined the response of the PARylation-deficient mutants to PARP inhibition. RADAR assays were performed in HCT116 cells expressing R338W TOP3B, R338W/E438A TOP3B, R338W/T488A TOP3B, or R338W/E438A/T488A TOP3B. PARP inhibition markedly increased TOP3Bcc accumulation in cells expressing R338W TOP3B but failed to further enhance TOP3Bccs in cells expressing the PARylation-deficient TOP3B mutants (**Figure 3H**). These results suggest that PARP1 activity and ADP-ribosylation of E438 and T488 act epistatically to regulate TOP3Bcc homeostatic levels. Together, these results identify E438 and T488 as major functional ADP-ribosylation sites on TOP3B and demonstrate that modification of these residues is required for efficient TOP3Bcc homeostasis.

### 4. PARylation recuits FEN1 to TOP3Bccs

To identify whether repair factors are recruited to TOP3Bcc following PARylation for their removal, we immunoprecipitated FLAG-tagged R338W TOP3B from HCT116 cells and analyzed associated proteins by LC-MS/MS. Among the proteins enriched following PARG inhibition, flap endonuclease 1 (FEN1), a structure-specific nuclease involved in nucleic acid processing and repair (32), showed a marked increase in association with TOP3B (**Figure 4A**).

**Figure 4.**
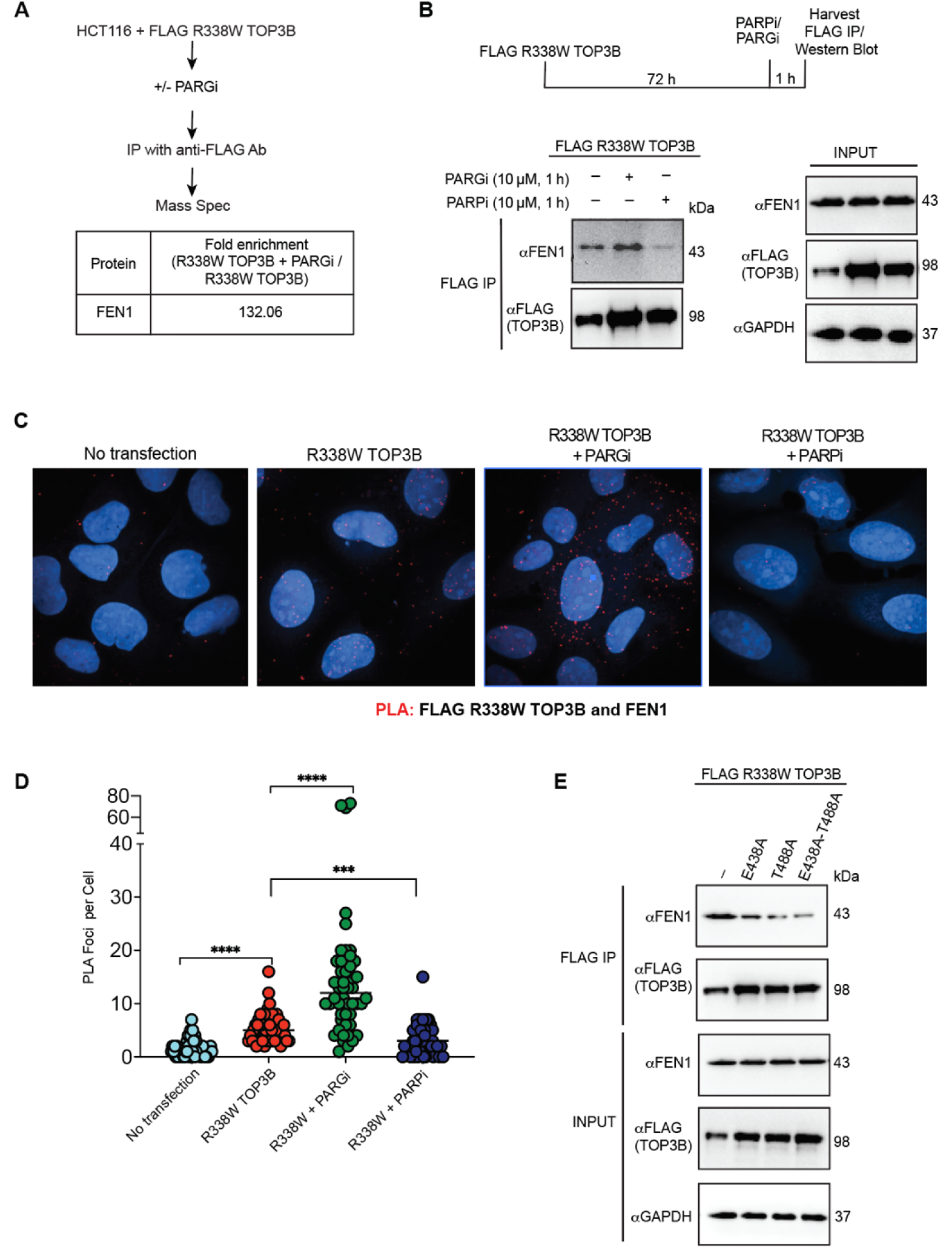
PARylation recruits FEN1 to TOP3B cleavage complexes. **A.** Proteomic analysis identifies enhanced association of FEN1 with TOP3B following PARG inhibition. FLAG R338W TOP3B was affinity-purified from HCT116 cells with or without PARGi treatment and analyzed by LC-MS/MS. Fold enrichment of FEN1 following PARG inhibition is shown. **B.** PARylation promotes the TOP3B-FEN1 interaction. HCT116 cells expressing FLAG R338W TOP3B were treated with PARGi or PARPi, followed by FLAG immunoprecipitation and immunoblotting for FEN1 and FLAG-TOP3B. Whole-cell lysates are shown as input controls. **C-D.** Proximity ligation assay confirms PARylation-dependent association of TOP3B with FEN1 in cells. C, representative PLA images detecting FLAG R338W TOP3B and endogenous FEN1 in untreated, PARGi-treated, and PARPi-treated HCT116 cells. D, quantification of PLA foci. n = 50 cells. Statistical significance was determined by two-tailed unpaired t test with Welch’s correction. ***P < 0.001; ****P < 0.0001. **E.** ADP-ribosylation-deficient TOP3B mutants show reduced interaction with FEN1. FLAG R338W TOP3B and the indicated R338W/E438A, R338W/T488A, or R338W/E438A/T488A TOP3B mutants were immunoprecipitated and analyzed for association with endogenous FEN1.

To validate the PARylation-dependent TOP3B-FEN1 interaction, we performed TOP3B immunoprecipitation using FLAG antibody in HCT116 cells expressing FLAG-tagged R338W TOP3B. Consistent with the proteomic results, PARG inhibition enhanced the association between TOP3B and FEN1, whereas PARP inhibition markedly reduced this interaction (**Figure 4B**). These findings indicate that TOP3B–FEN1 association is promoted by PARylation.

To confirm this TOP3Bcc-FEN1 interaction *in situ*, we performed proximity ligation assays (PLA) in HCT116 cells expressing FLAG-tagged R338W TOP3B with or without PARP or PARG inhibition. FEN1–FLAG-TOP3B PLA foci were observed both in the nucleus and the cytoplasm in the transfected cells, supporting the specificity of the signal and confirming the proximity of FEN1 and TOP3B in cells. PARG inhibition further increased the FEN1-TOP3B PLA foci whereas PARP inhibition significantly reduced the number of the foci (**Figures 4C and 4D**). These results support a PARylation-dependent interaction between TOP3B and FEN1.

Finally, we examined whether the major TOP3B ADP-ribosylation sites identified above are required for FEN1 association. FLAG immunoprecipitation followed by immunoblotting was performed in HCT116 cells expressing FLAG-tagged R338W TOP3B or the PARylation-deficient R338W/E438A, R338W/T488A, and R338W/E438A/T488A mutants. All three mutants showed reduced association with FEN1 compared with R338W TOP3B, with the R338W/E438A/T488A mutant displaying the strongest defect (**Figure 4E**). Together, these results demonstrate that TOP3B PARylation at E438 and T488 promotes recruitment of FEN1 to TOP3B.

### 5. PARylation promotes excision of TOP3Bccs by FEN1

To determine whether PARylation-dependent recruitment of FEN1 removes TOP3Bccs, we depleted FEN1 in HCT116 cells expressing FLAG-tagged R338W TOP3B. RADAR assays showed that FEN1 depletion increased TOP3Bccs (**Figure 5A**). Consistent with our earlier results (see Figure 1), PARP inhibition also increased TOP3Bcc levels. Combined FEN1 depletion and PARP inhibition did not further increase TOP3Bcc levels beyond that observed with PARP inhibition alone (**Figure 5A**). This suggests an epistatic relationship between FEN1 and PARP1 in TOP3Bcc removal.

**Figure 5.**
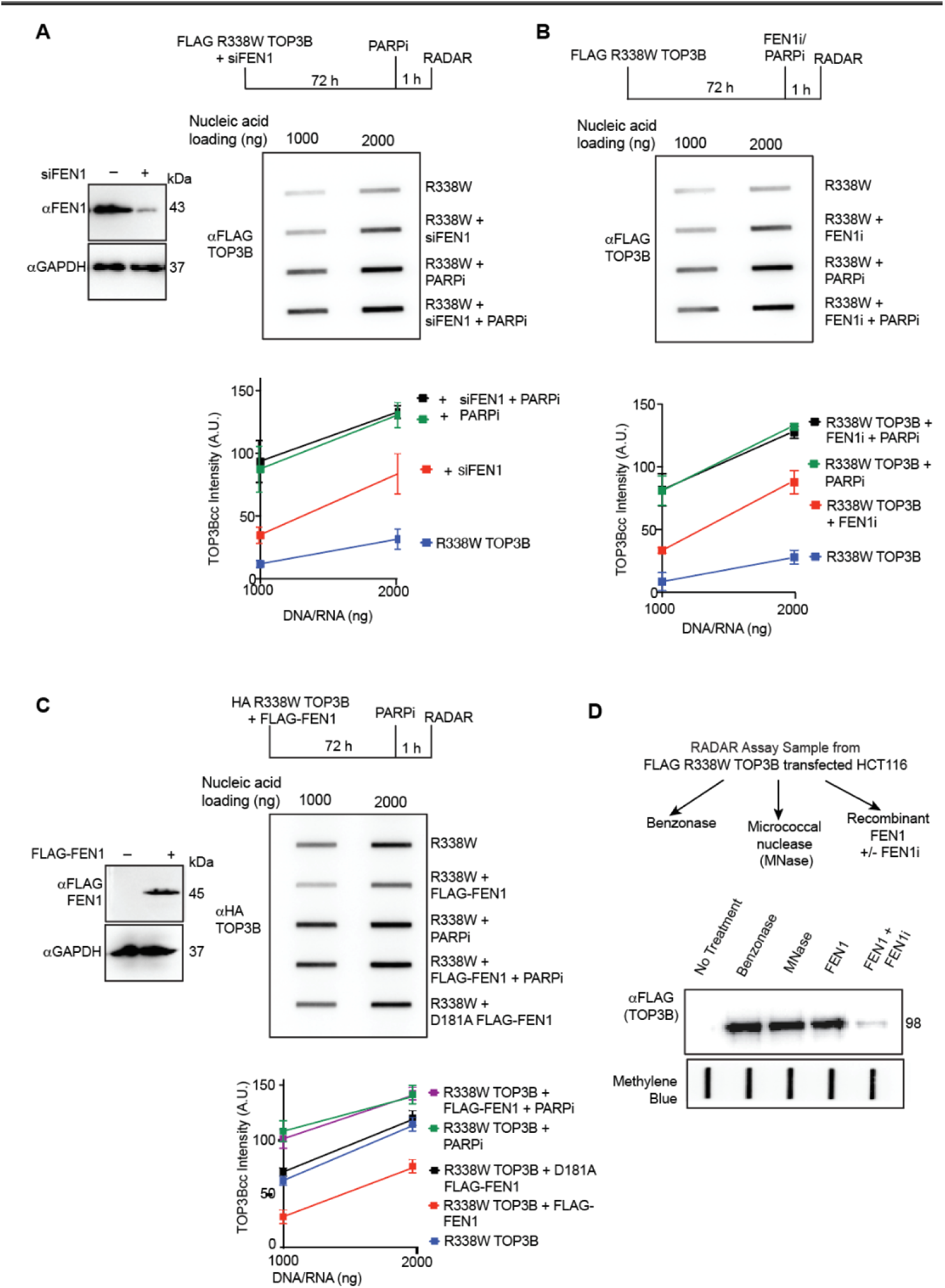
FEN1 removes TOP3B cleavage complexes in a PARP1-dependent manner. **A.** FEN1 depletion and PARP inhibition are epistatic for TOP3Bcc accumulation. HCT116 cells expressing FLAG R338W TOP3B were transfected with control or FEN1 siRNA and treated with PARPi as indicated. FEN1 depletion was confirmed by immunoblotting, and TOP3Bcc levels were measured by RADAR slot blotting. **B.** Pharmacological inhibition of FEN1 does not further increase TOP3Bcc levels following PARP inhibition. HCT116 cells expressing FLAG R338W TOP3B were treated with PARPi, FEN1 inhibitor (10 uM, 1h), or both, and TOP3Bccs were quantified by RADAR slot blotting. **C.** FEN1 nuclease activity promotes TOP3Bcc removal in a PARP1-dependent manner. HCT116 cells expressing HA-tagged R338W TOP3B were co-transfected with wild-type FLAG-tagged FEN1 or nuclease-deficient FEN1 D181A and treated with PARPi as indicated. FEN1 expression and TOP3Bcc levels were analyzed by immunoblotting and RADAR slot blotting, respectively. **D.** Recombinant FEN1 releases TOP3B from isolated cellular TOP3Bccs. RADAR samples containing FLAG R338W TOP3B were incubated with recombinant FEN1, FEN1 plus FEN1 inhibitor, benzonase, or micrococcal nuclease. Released TOP3B was detected by immunoblotting with anti-FLAG antibody. n = 3 independent experiments.

To confirm the role of FEN1 in TOP3Bcc removal, we treated HCT116 cells expressing R338W TOP3B with the FEN1 inhibitor FEN1-IN-4 (33). Inhibition of either FEN1 or PARP increased TOP3Bccs (**Figure 5B**). Yet, combined inhibition of FEN1 and PARP did not produce further increase in TOP3Bcc levels (**Figure 5B**). These results support the conclusion that FEN1 and PARP1 function in the same repair pathway for TOP3Bcc removal.

To establish whether FEN1’s nuclease activity is required for TOP3Bcc resolution, HCT116 cells expressing HA-tagged R338W TOP3B were co-transfected with FLAG-tagged wild-type FEN1 or the nuclease-deficient FEN1 D181A mutant with or without PARP inhibitor treatment. Upregulation of wild-type FEN1 reduced cellular TOP3Bcc levels, whereas the D181A mutant failed to remove cellular TOP3Bccs (**Figure 5C**). These results indicate that FEN1 nuclease activity is required for efficient TOP3Bcc removal. Notably, wild-type FEN1 failed to remove cellular TOP3Bccs in the presence of PARP inhibitor, supporting the functional coupling between PARylation and FEN1-mediated TOP3Bcc repair (**Figure 5C**).

We also assessed whether FEN1 can excise T*OP3Bccs in vitro*. To do so, we isolated RADAR samples from HCT116 cells expressing FLAG-tagged R338W TOP3B and incubated with recombinant FEN1 protein, recombinant FEN1 protein pre-incubated with the FEN1 inhibitor, benzonase, or micrococcal nuclease before SDS-PAGE analysis. Recombinant FEN1 released TOP3B from nucleic acid-linked complexes, allowing detection of free TOP3B by SDS-PAGE, whereas inhibition of FEN1 nuclease activity with FEN1-IN-4 prevented this effect (**Figure 5D**). As expected, benzonase and micrococcal nuclease efficiently released TOP3B and served as positive controls (**Figure 5D**). Together, these results demonstrate that FEN1 excises TOP3Bccs through its nuclease activity and that efficient FEN1-mediated TOP3Bcc removal depends on PARP1 activity. These findings support a model in which PARylation facilitates FEN1 recruitment to TOP3Bccs, enabling their subsequent nucleolytic excision.

### 6. Persistent PARylation recruits USP10 and promotes TOP3Bcc deubiquitylation and stabilization

We next investigated why persistent PARylation induced by PARG inhibition leads to TOP3Bcc accumulation. Because ubiquitylation is a major determinant of TOP3Bcc proteasomal processing, we tested whether persistent PARylation alters the ubiquitylation state of TOP3Bccs. To examine the relationship between TOP3Bcc PARylation, ubiquitylation, and stability, we performed the modified RADAR assay (34) in cells expressing FLAG-tagged R338W TOP3B following PARG inhibition for the indicated times.

Consistent with the experiments shown in Figure 1, PARG inhibition caused a progressive increase in TOP3Bcc levels (**Figure 6A**). As expected, PARG inhibition also led to a time-dependent increase in PARylated TOP3Bcc species, which became readily detectable within 30 min. In contrast, TOP3Bcc ubiquitylation progressively decreased following PARG inhibition (**Figure 6A**). These reciprocal changes suggest that persistent PARylation may stabilize TOP3Bccs by antagonizing their ubiquitylation and subsequent proteasomal processing.

**Figure 6.**
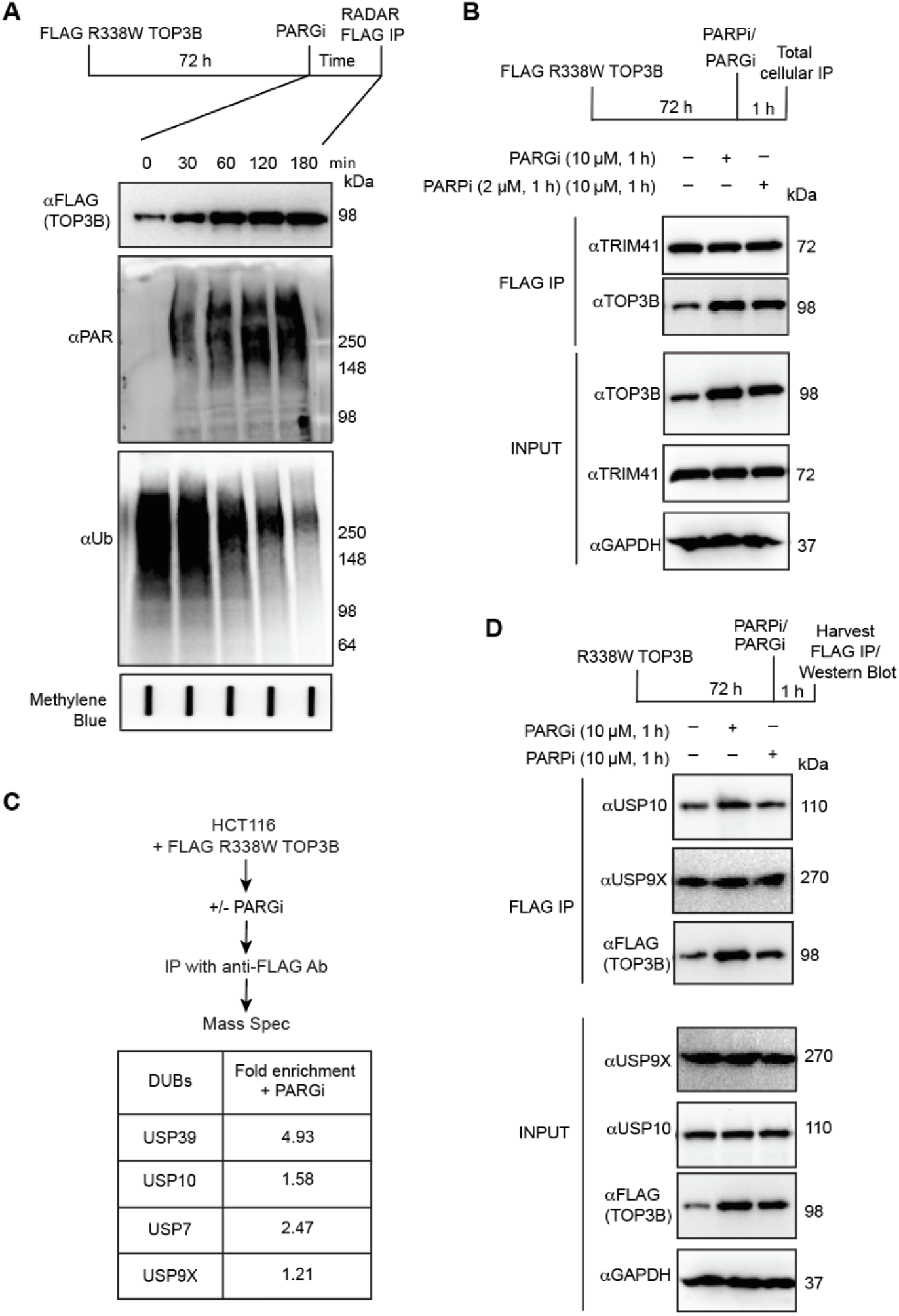
Persistent PARylation recruits USP10 and suppresses TOP3Bcc ubiquitylation. **A.** Persistent PARylation following PARG inhibition is associated with reduced TOP3Bcc ubiquitylation. HCT116 cells expressing FLAG R338W TOP3B were treated with PARGi (10 uM) for the indicated time periods. TOP3Bccs isolated by RADAR were immunoprecipitated with anti-FLAG antibody and analyzed by immunoblotting for PAR, ubiquitin, and FLAG-tagged TOP3B. **B.** Effect of PARG inhibition on the interaction of TOP3B with TRIM41 protein. FLAG R338W TOP3B was immunoprecipitated from control, PARPi or PARGi-treated HCT116 cells and associated TRIM41 protein was examined by immunoblotting. **C.** Proteomic analysis identifies USP10 as a PARGi-enhanced TOP3B-interacting deubiquitylase. FLAG R338W TOP3B was affinity-purified from control or PARGi-treated HCT116 cells and analyzed by LC-MS/MS. Fold enrichment of the indicated deubiquitylases following PARG inhibition is shown. **D.** PARylation promotes the TOP3B-USP10 interaction. HCT116 cells expressing FLAG-TOP3B R338W were treated with PARGi or PARPi, followed by FLAG immunoprecipitation and immunoblotting for USP10, USP9X, and FLAG-TOP3B. Whole-cell lysates are shown as input controls.

Because the E3 ubiquitin ligase TRIM41 has been shown to ubiquitylate TOP3Bccs and promote the proteasomal processing of TOP3Bccs (12), we asked whether PARylation interferes with the TOP3B–TRIM41 interaction. FLAG immunoprecipitation in HCT116 cells expressing FLAG-tagged R338W TOP3B showed that TOP3B interacts with TRIM41 independently of PARP or PARG activity (**Figure 6B**).

To test whether the reduction in TOP3Bcc ubiquitylation is related to the recruitment of a deubiquitylating enzyme (DUB), we performed FLAG immunoprecipitation followed by LC-MS/MS in cells expressing FLAG-tagged R338W TOP3B. PARG inhibition promoted interaction of TOP3B with several DUBs, including USP10 and USP9X (**Figure 6C**). We focused on USP9X and USP10 because USP9X is a previously reported DUB for TOP3B (16) and USP10 is known to preferentially interact with PARylated substrates (35,36). TOP3B FLAG IP-Western blotting showed that PARG inhibition enhanced the TOP3B-USP10 interaction, whereas PARP inhibition reduced this interaction (**Figure 6D**). Yet, neither PARP nor PARG inhibition appreciably altered the interaction between TOP3B and USP9X. These findings indicate that persistent PARylation promotes the recruitment of USP10 to TOP3Bccs.

We next examined whether USP10 regulates the ubiquitylation and stability of persistently PARylated TOP3Bccs. HCT116 cells were co-transfected with FLAG-tagged R338W TOP3B and USP10 siRNA and analyzed by the modified RADAR assay (34) (**Figure 7A**). Downregulation of USP10 by siRNA (**Figure 7B**) resulted in enhanced TOP3Bcc ubiquitylation and reduced TOP3Bcc signal (**Figure 7C**), with these effects being particularly pronounced following PARG inhibition (**Figure 7C**). These results imply that USP10 contributes to the deubiquitylation and stabilization of persistently PARylated TOP3Bccs. Conversely, ectopic expression of USP10 (**Figure 7E**) resulted in deubiquitylation and stabilization of TOP**3**Bccs (**Fi**g**ure 7F**) both in the absence and presence of PARG inhibition. Notably, combination of PARGi and USP10 overexpression further enhanced TOP3Bcc deubiquitylation and stabilization (**Figure 7F**). Together, these results support a model in which PARylation, if not reversed promptly, recruits USP10 to TOP3Bccs, leading to their deubiquitylation and stabilization. Thus, whereas transient PARylation promotes TOP3Bcc repair through FEN1 recruitment, persistent PARylation shifts TOP3Bcc processing toward USP10-dependent stabilization.

**Figure 7.**
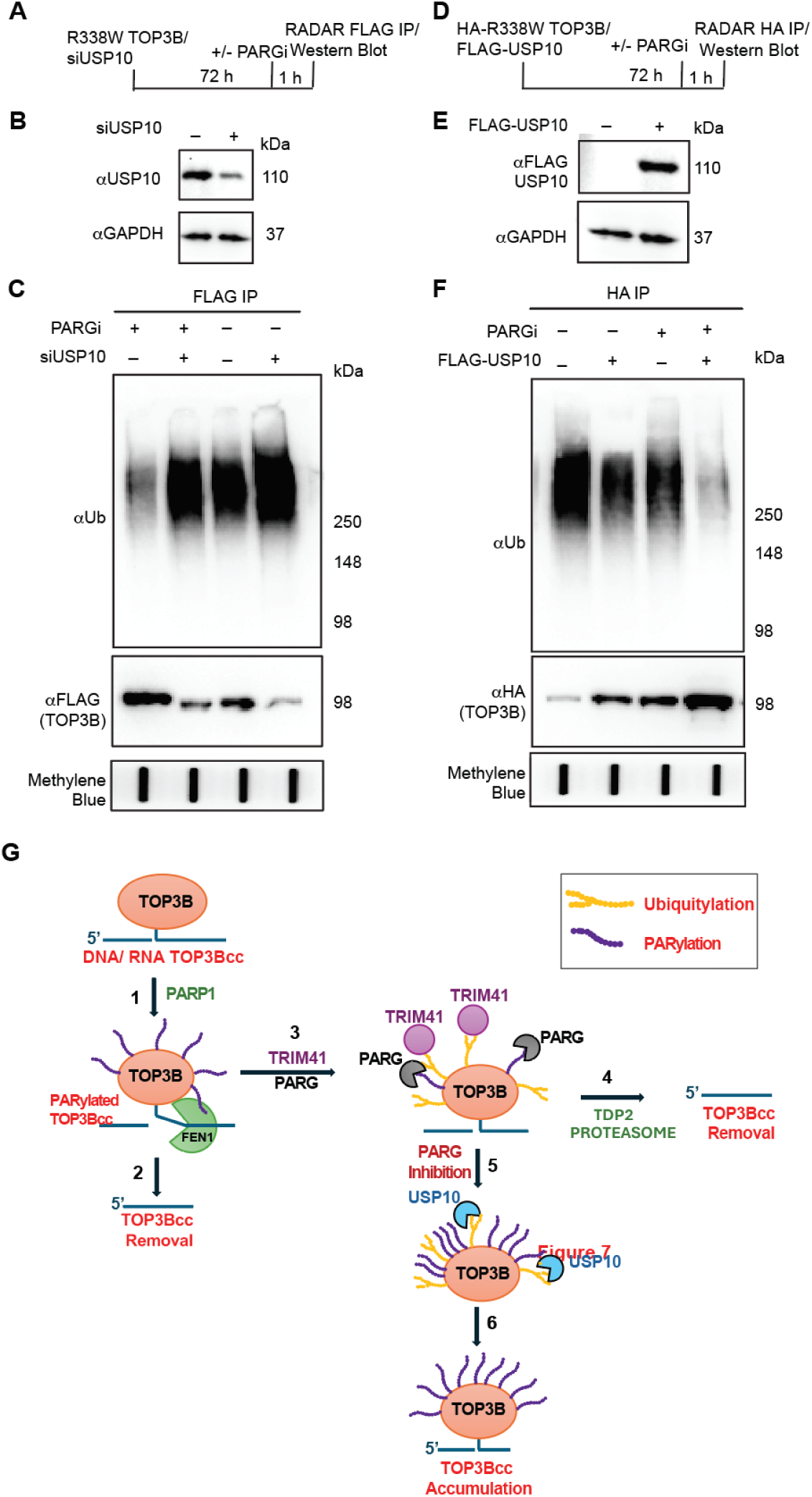
USP10 promotes deubiquitylation and stabilization of PARylated TOP3Bccs and proposed model of PARylation-dependent cellular TOP3Bcc homeostasis. **A.** Experimental design for the modified RADAR assay determining the effect of USP10 depletion on ubiquitylation and stability of persistently PARylated TOP3Bccs. **B.** Western blotting confirming USP10 depletion in HCT116 cells. **C.** HCT116 cells expressing FLAG R338W TOP3B were transfected with control or USP10 siRNA and treated with PARGi as indicated. TOP3Bccs isolated by RADAR were immunoprecipitated with anti-FLAG antibody and analyzed by immunoblotting for ubiquitin and FLAG-tagged TOP3B. **D.** Experimental design for the modified RADAR assay determining the effect of USP10 upregulation on ubiquitylation and stability of persistently PARylated TOP3Bccs. **E.** Western blotting confirming FLAG-USP10 overexpression. **F.** HCT116 cells expressing HA-tagged R338W TOP3B were co-transfected with FLAG-USP10 and treated with PARGi as indicated. TOP3Bccs isolated by RADAR were immunoprecipitated with anti-HA antibody and analyzed by immunoblotting for ubiquitin and HA-tagged TOP3B. **G.** Model depicting PARylation dependent regulation of cellular TOP3Bcc homeostasis. TOP3Bcc formation activates PARP1 (1) which rapidly and transiently PARylates TOP3Bccs formed both in DNA and RNA. PARylation recruits structure-specific endonuclease FEN1 for TOP3Bcc repair (2). Later, PARG protein removes TOP3Bcc PARylation while, in parallel, the E3 ubiquitin ligase TRIM41 ubiquitylates TOP3Bcc (3). Ubiquitylation targets TOP3Bcc for proteasomal processing and finally TDP2 hydrolyzes the exposed phosphotyrosyl bond to excise the TOP3B polypeptide remaining attached to the 5’-end of DNA and RNA molecules (4). Inhibition of PARG activity causes PAR chains to accumulate on TOP3Bccs (5), recruits the DUB USP10 (5), which deubiquitylates TOP3Bccs and prevents proteasomal degradation of DNA- and RNA-linked TOP3Bccs (6).

## DISCUSSION

Human topoisomerase IIIβ (TOP3B) is the only human topoisomerase that acts on both DNA and RNA (12–14). TOP3Bcc have been shown to process R-loops, hypernegatively supercoiled DNA, RNA knots and catenanes (37). It has also been implicated as a prognostic biomarker and potential therapeutic target in many types of cancers (38–40) and its alterations cause neurological disorders (17,41,42). We have recently identified first-in-class human TOP3B inhibitors that trap TOP3Bccs by blocking their reversal (19). Resulting abortive TOP3Bccs disrupt nucleic acid transactions, induce nucleic acid damage and lead to cell death (12,16,19). As with other topoisomerase-targeting agents, the therapeutic efficacy of TOP3B inhibitors is likely to be influenced by cellular pathways that recognize and repair drug-stabilized cleavage complexes. Understanding how cells process DNA- and RNA-linked TOP3Bccs is therefore important both for defining TOP3B biology and for identifying mechanisms that may modulate sensitivity or resistance to TOP3B-targeting agents (12).PARylation, a chemical post-translational modification, has previously been implicated in the processing of trapped TOP1ccs, TOP2ccs and TOP3Accs (34,43–46). In response to topoisomerase trapping, PARP1 rapidly catalyzes transient PARylation of itself and trapped topoisomerase complexes, thereby facilitating the recruitment of downstream repair factors. For example, PARylation promotes recruitment of TDP1 to TOP1ccs (47), while PARP inhibition increases the accumulation of these lesions (34,43,44). Yet, whether PARylation regulates TOP3Bccs, particularly RNA-linked TOP3Bccs, has not been established.

In the present study, using an unbiased chemical screen, we identified PARP inhibitors, including olaparib, rucaparib and niraparib, as synthetic lethal with the TOP3B-trapping agent NSC690634. This observation was consistent with our previous finding that TOP3Bcc accumulation enhances the interaction between TOP3B and PARP1 (16), and prompted us to interrogate whether PARylation directly regulates TOP3Bcc repair. Our results reveal a dynamic PARylation-dependent pathway that controls the homeostasis of both DNA- and RNA-linked TOP3Bccs (as summarized in **Figure 7G**). We show that TOP3Bccs are transiently PARylated and only readily detectable when PAR turnover is blocked by PARG inhibition. Notably, inhibition of either PAR synthesis or PAR removal increase TOP3Bcc accumulation, indicating that efficient TOP3Bcc resolution requires dynamic PARylation rather than simply the presence of PAR. We identify the amino acid residues E438 and T488 as major functional ADP-ribosylation sites on TOP3B and show that mutating these residues markedly reduce TOP3B PARylation and increase TOP3Bcc accumulation. The role of PARylation in the repair of RNA-linked TOP3Bccs is particularly notable. PARP1 and PARP2 are best characterized as sensors of DNA lesions and can be activated by single-stranded DNA, DNA breaks, hairpins, cruciform structures, and R-loops (48–51). Also, PARP enzymes have been implicated in multiple steps of RNA metabolism—from transcription, splicing, polyadenylation, export, translation of mRNA, RNA stress granule dynamics to RNA degradation (50,52–62). PARP proteins have been shown to physically bind to RNA molecules (63,64) and transcriptome-wide studies have identified extensive PARP1–RNA interactions (65–68). PARP-1 catalytic activity is stimulated by various forms of RNA (69–71). For example, binding of small nucleolar RNAs (snoRNAs) to the “DNA-binding” domain of PARP-1 activates PARP1, which in turn PARylates the RNA helicase DDX21 (65,72). To our knowledge, our data provide the first evidence that RNA-linked TOP3Bccs are PARylated and repaired by a PARP1-dependent repair pathway. These findings extend the biological scope of PARylation-mediated topoisomerase repair beyond DNA and raise the possibility that PARP-dependent signaling participates directly in the surveillance of protein-linked RNA lesions.

Mechanistically, our study identifies FEN1 as a downstream effector of PARylation-dependent TOP3Bcc repair. FEN1 plays pivotal roles in Okazaki-fragment maturation and long-patch base-excision repair, where it removes 5′-flap structures (73–76). It has also been implicated in the processing of TOP2ccs and TOP3Accs containing 5′-phosphotyrosyl linkages (43,44,77). FEN1 can also process RNA-containing flap structures, including RNA within R-loops (76,78,79). Here, we find that FEN1 also removes TOP3Bccs from the 5′-ends of nucleic acids in a PARylation-dependent manner. Our PLA experiments show that PARylation promotes the interactions between FEN1 and TOP3B both in the nucleus and the cytoplasm. As cytoplasm primarily harbors RNA-linked TOP3Bccs, this observation is consistent with the possibility that FEN1 participates in processing both DNA- and RNA-associated TOP3B lesions. The mechanistic details by which FEN1 excises RNA-linked TOP3Bccs warrant further investigations.

Our findings also reveal an unexpected consequence of persistent PARylation. We observed a reciprocal relationship between PARylation and ubiquitylation following PARG inhibition. Persistent PARylation of TOP3Bccs increased with time while TOP3Bcc ubiquitylation progressively decreased. Rather than preventing forward ubiquitylation, persistent PARylation appears to promote active deubiquitylation of TOP3Bccs. Proteomic analysis identified several DUBs associated with TOP3B following PARG inhibition. Among these, USP10 showed a clear PARylation-dependent interaction. PARG inhibition enhanced TOP3B–USP10 association, whereas PARP inhibition markedly reduced it. Functionally, USP10 depletion increased TOP3Bcc ubiquitylation and reduced TOP3Bcc abundance, whereas USP10 overexpression decreased ubiquitylation and stabilized TOP3Bccs, particularly under conditions of PARG inhibition. These results support a model in which persistent PARylation recruits USP10 to deubiquitylate and stabilize TOP3Bccs, presumably by preventing proteasomal degradation (**Figure 7G**).

In summary, our study reveals a role for PARylation as a dynamic molecular switch in TOP3Bcc homeostasis. Transient PARylation promotes the recruitment of FEN1 and facilitates TOP3Bcc excision, whereas persistent PARylation redirects TOP3Bcc processing toward USP10-dependent deubiquitylation and stabilization under conditions of PARG inhibition (**Figure 7G**). Our findings not only reveal the different pathways involved in the removal of potentially toxic TOP3Bccs, but also may have therapeutic implications. PARP inhibitors are widely used as anticancer agents and their impact on TOP3Bccs may need to be considered. Similarly, PARG inhibitors are being developed as anticancer agents and their inhibitory impact on the repair of TOP3Bccs may be a source of genotoxic lesions. FEN1 inhibition may provide a third strategy to interfere with this TOP3Bcc repair pathway. The therapeutic potential of combining TOP3B inhibitors with PARP or FEN1 inhibitors will require direct validation across additional cancer models, in vivo efficacy s*tudies,* and assessment of therapeutic index. More broadly, defining the pathways that govern TOP3Bcc repair should help clarify which tumors are most likely to depend on TOP3B and which repair vulnerabilities can be exploited to enhance responses to TOP3B-targeting therapies.

## MATERIALS AND METHODS

### 1. Cell lines

HCT116 cells (Developmental Therapeutics Program, National Cancer Institute) were grown in Dulbecco’s modified Eagle’s medium (DMEM, ThermoFisher Scientific, Cat# 11965-092) supplemented with 10% Fetal Bovine Serum (Gemini, West Sacramento, CA, 100-106) and 1% penicillin-streptomycin (ThermoFisher Scientific, 15140122) at 37°C in a humidified 5% CO2 incubator. DMS-114 cell line was purchased from the ATCC and maintained in culture in RPMI 1640 medium supplemented with 10% FBS, penicillin/streptomycin 1×, and GlutaMax 1×. Cell lines were tested regularly for mycoplasma contamination (Lonza, LT07-318) and were confirmed to be free of mycoplasma contamination. HCT116 and DMS114 cell line authentications were carried out using short tandem repeat (STR) profiling analysis.

### 2. Chemicals

The chemicals used are as follows: PDD 00017273 (PARGi, Tocris Bioscience, Cat# 5952); Olaparib (PARPi, Selleck, Cat# S1060), FEN1-IN-4 (FEN1i, Selleck, Cat# S3397), Methylene Blue Hydrate (SigmaAldrich, Cat# M4159-25G)

### 3. Mammalian expression constructs and transient expression in mammalian cells

The following expression plasmids were used: pShuttle--FLAG--FEN1, Addgene, 35027; Myc-FLAG-tagged R338W TOP3B(12); Human TOP3B subcloned into pcDNA3-HA with BamHI and EcoRI sites(12); Myc-FLAG-tagged USP10, OriGene, RC200835; Plasmids were transfected in HCT116 cells using Lipofectamine 3000 Reagent (CAT#: L3000015, ThermoFisher Scientific) according to the manufacturer’s protocol for 48–72 h.

### 4. Antibodies

The following antibodies were used: Mouse monoclonal Anti-PAR Antibody , R&D Systems, Inc. a Bio-Techne Brand, CAT#: 4335-MC-100; Mouse monoclonal anti-FLAG M2, Millipore Sigma, St. Louis, MO, CAT#: F1804); Rabbit monoclonal anti-GAPDH, Cell Signaling Technology, Danvers, MA, CAT#: 2118 S; Rabbit monoclonal anti-HA, Cell Signaling Technology, Danvers, MA, CAT#: 3724 S); anti–double-stranded DNA (dsDNA), mouse monoclonal, Abcam, ab27156; Rabbit monoclonal anti-FEN1, Cell Signaling Technology, 82354S; Rabbit monoclonal anti-TOP3B antibody, Abcam, Waltham, MA, CAT#: ab183520; Rabbit monoclonal anti-USP9X antibody, Cell Signaling Technology, Danvers, MA, CAT#: 14898S; Rabbit monoclonal anti-USP10 antibody, Cell Signaling Technology, Danvers, MA, CAT#: 8501S; Mouse monoclonal Anti-Ubiquitin Antibody (P4D1), Santa Cruz Biotechnology Cat# sc-8017, RRID:AB_628423; Sheep Anti-Mouse IgG ECL Antibody, HRP Conjugated, GE Healthcare,Cat # NA9310-1ml, RRID:AB_772193; Donkey Anti-Rabbit IgG ECL Antibody, HRP Conjugated, GE Healthcare, Cat# NA9340-1ml, RRID:AB_772191.

### 5. siRNA Transfection

The following siRNAs were used: ON-TARGETplus SMARTpool Human FEN1 siRNA (CAT#: L-010344-00-0005); ON-TARGETplus SMARTpool Human USP10 siRNA (CAT#: L-006062-00-0005); ON-TARGETplus Non-targeting Control Pool (CAT#: D-001810-10-05). All siRNAs were used at a final concentration of 25 nM and transfected using Lipofectamine® RNAiMAX transfection reagent (CAT#: 13778150, ThermoFisher Scientific) following the manufacturer’s protocol for 48–72 h.

### 6. Recombinant proteins

Human recombinant FEN1 is a gift from R. Prasad at NIEHS. Recombinant human PARP1 was expressed in *Escherichia coli* BL21(DE3) cells. Clarified cell lysate was applied to a nickel-affinity column, and bound protein was eluted with 300 mM imidazole. PARP1 was then purified sequentially on heparin and SP columns, each using a linear gradient of 50 mM-1 M NaCl in 50 mM Tris-HCl (pH 7.0), 1 mM EDTA, 1 mM DTT, and 10% glycerol. Fractions containing PARP1 were pooled, concentrated, and loaded onto a Superdex 200 10/300 column equilibrated in 25 mM Tris-HCl (pH 7.5), 150 mM NaCl, 1 mM EDTA, 1 mM DTT, and 10% glycerol. Protein purity was assessed by SDS-PAGE. Purified PARP1 was concentrated, aliquoted, and flash-frozen in liquid nitrogen.

### 7. High-throughput chemical screening

High-throughput cell viability assays were performed in DMS114 small cell lung cancer (SCLC) cells in a 384-well format by the Target Validation and Screening (TVS) Service of the Center of Biomolecular Therapeutics (CBT). This study used the FDA-Approved Anticancer Drug Library (Selleck Chemicals), a collection of 1728 approved drugs with potential anticancer activity, to identify compounds that alter cellular responses to TOP3B trapping. A 1 mM dilution of the drug library was prepared in 384-well format using a Biomek i7 automated workstation (Beckman-Coulter) for this study.

DMS114 cells were plated at 500 cells/well in a volume of 20 µl phenol-red free media in Falcon™ Optilux™ 384-well cell culture plates (Fisher Scientific). After recovering for 24 hours at 37 °C and 5% CO_2_, the cells were treated with 1 µM doses of library compounds from 1 mM stock plates using the Biomek i7 with a pin tool attachment (V&P Scientific) containing 20 nL slotted pins. The cells were incubated for a further 4 hours before being dosed with or without an IC_20_ dose of TOP3B poison NSC690634 (final concentration: 2.5 nM). The cells were then returned to the incubator for 72 hours. Cell viability analysis was performed using CellTiter-Glo 2.0 kit (Promega) following the manufacturer’s protocol. A ClarioStar Plus plate reader (BMG Labtech) was used to measure luminescence.

Raw luminescence was normalized to the DMSO controls within a 384-well plate to give the viability of the drug-treated cell populations. However, as the control luminescence showed a distinct edge effect (loss of over 20% signal in the outer wells), the drug wells in the top and bottom rows (A and P, respectively) were normalized to the outer DMSO control wells (columns 1 and 24), while the inner drug wells in rows B to O were standardized to the inner DMSO control wells (columns 2 and 23). For plates that contained aqueous-based drugs, the luminescence values were normalized to the appropriate aqueous controls (i.e., inner versus outer).

For each compound, the change in normalized viability was calculated as ΔVᵢ = V₍alone,ᵢ₎ − V₍combination,ᵢ₎. To display greater apparent sensitization as a more negative score, viability differences were standardized as Zᵢ = (μΔV − ΔVᵢ) / σΔV, where μΔV and σΔV are the mean and standard deviation of the analyzed compound-level viability differences, respectively. Compounds were ranked by Z-score, with more negative values indicating greater apparent sensitization to NSC690634. These single-dose scores assess differential viability and do not establish pharmacological synergy.

### 8. Proximity ligation assay

Duolink PLA fluorescence assay (Sigma--Aldrich, catalog no. DUO92101) was performed following the manufacturer’s instructions. In brief, HCT116 cells transiently transfected with FLAG-tagged R338W TOP3B were seeded on coverslips overnight. Cells were washed with 1 × phosphate--buffered saline (PBS) and fixed for 15 min at 4°C in 4% paraformaldehyde (PFA) in PBS and permeabilized with 0.25% Triton X--100 in PBS for 15 min at 4°C. The coverslips were blocked with Duolink blocking solution and incubated with indicated antibodies in the Duolink antibody diluent overnight, followed by incubation with PLUS and MINUS PLA probes, ligation, and amplification. The coverslips were then washed and mounted using mounting medium with 4 ′,6-­ diamidino-­ 2-­ phenylindole (DAPI). Images were captured on a wide-­ field microscope, processed using ImageJ, and analyzed using Imaris.

### 9. Western blotting

To prepare whole cell lysates for western blotting, cells were resuspended with RIPA buffer (150 mM NaCl, 1% NP-40, 0.5% Sodium deoxycholate, 0.1% SDS, 50 mM Tris pH 7.5, 1 mM DTT and cOmplete Mini, EDTA-free protease inhibitor cocktail (Roche, Cat# 11836170001). After thorough mixing, samples were agitated at 4 °C for 30 min, sonicated for 30 s with 40% pulse (3-5 times), centrifuged at 15,000 × g at 4 °C for 15 min, and supernatant was collected.

For detection of total cellular TOP3B (both free and nucleic acid bound), cells were washed with PBS and incubated on a shaker with IP lysis buffer (50 mM Tris-HCl pH 7.4, 150 mM NaCl, 1 mM EDTA, 1% NP-40, 0.2% Triton X-100, 5% glycerol, 1 mM DTT, 20 mM N-ethylmaleimide (Millipore Sigma, Cat# E3876-25G) and protease inhibitor cocktail supplemented with 2 μl benzonase (250 units/μl, Sigma-Aldrich, CAT#: E8263) followed by 30 min incubation in ice. After brief sonication (40% power for 30 s pulse and 1 min rest 4 times), samples were centrifuged at 15,000×g at 4 °C for 30 min and the supernatant was collected.

Lysed samples were mixed with tris-glycine SDS sample buffer (Novex, LC2676) and loaded onto Novex tris-glycine gels (Novex). Blotted membranes were blocked with 5% non-fat dry milk in PBS with 0.1% Tween-20 (PBST). The primary antibodies were diluted in 5% milk in PBST, and secondary antibodies were diluted (1:10000) in 5% non-fat milk in PBST and signal was detected by ECL chemiluminescence reaction (SuperSignal™ West Femto Maximum Sensitivity Substrate, Thermo Scientific, Waltham, MA, Cat# 34095).

### 10. Immunoprecipitation (IP)

Cells were washed with PBS and incubated on a shaker with IP lysis buffer (50 mM Tris-HCl pH 7.4, 150 mM NaCl, 1 mM EDTA, 1% NP-40, 0.2% Triton X-100, 5% glycerol, 1 mM DTT, 20 mM N-ethylmaleimide and protease inhibitor cocktail) supplemented with 2 μl benzonase (250 units/μl, Sigma-Aldrich, CAT#: E8263) followed by 30 minutes incubation in ice. After brief sonication (40% power for 30 s pulse and 1 min rest 3 times), samples were centrifuged at 15,000 X g at 4 °C for 30 min and the supernatant was collected and an aliquot (20 μl) of the supernatant was saved as input. The rest of the supernatant was diluted in IP lysis buffer containing desired antibody (3 μg/ tube) and rotated overnight at 4 °C. Next day, 30 μl Protein A/G magnetic beads (ThermoFisher Scientific Cat#: 26162) were added and incubated with the lysates for another 4 h. After magnetic separation, beads were washed with RIPA buffer 2 times. Finally, beads were resuspended in Tris-Glycine SDS Sample Buffer (2X) (ThermoFisher Scientific Novex™; Cat#: LC2676) for SDS-PAGE and immunoblotted with different antibodies as indicated or further processed for MS analysis.

### 11. RADAR Assay and detection of DNA and RNA TOP3Bccs

RADAR assay was performed for detection of TOP3Bccs as described previously(12,16). HA or FLAG-tagged R338W TOP3B transfected or NSC690634 treated HCT116 cells (1 × 10^6^) treated were harvested after 48–72 h, washed with PBS, and lysed by adding 1 mL DNAzol (ThermoFisher Scientific, CAT#:10503027). Nucleic acids were precipitated following addition of 0.5 mL of 100% ethanol, incubation at −20 °C for 5 min and centrifugation (12,000 × g for 10 min). Precipitates were washed twice in 75% ethanol, resuspended in 200 μl TE buffer, heated at 65 °C for 15 min, followed by shearing with sonication (40% power for 15 s pulse and 30 s rest 5 times). Samples were centrifuged at 15,000 rpm for 5 min and the supernatant containing nucleic acids with covalently bound proteins were collected. Nucleic acid-containing protein adducts were quantitated, slot-blotted and TOP3Bccs were detected with either rabbit monoclonal Anti-TOP3B antibody [EP7779]—C-terminal (Abcam, CAT#: ab183520) or mouse monoclonal anti-FLAG M2 antibody (Millipore Sigma, St. Louis, MO, CAT#: F1804) or rabbit Anti-HA-Tag Monoclonal Antibody, Unconjugated, Clone C29F4 (Cell Signaling Technology, CAT#: 3724).

For detection of DNA and RNA TOP3Bccs, 10 μg RADAR assay samples were digested either with excess RNase A (200 μg/mL; ThermoFisher Scientific Cat# EN0531) and RNase T1 (200 units/ml; ThermoFisher Scientific, Cat# EN0542) mix, or with DNase 1 (10 units; Invitrogen™ TURBO™ DNase (2 U/µL), ThermoFisher Scientific, Cat# AM2238) or Benzonase (250 units; Sigma-Aldrich, Cat# E8263). Samples were ethanol-precipitated, resuspended, quantitated, slot-blotted and TOP3Bccs were detected with Anti-TOP3B antibody [EP7779]—C-terminal (Abcam, CAT#: ab183520); or mouse monoclonal anti-FLAG M2 antibody (Millipore Sigma, St. Louis, MO, CAT#: F1804) or rabbit Anti-HA-Tag Monoclonal Antibody, Unconjugated, Clone C29F4 (Cell Signaling Technology, CAT#: 3724)(12,16,19).

### 12. Immunoprecipitation and enrichment of RADAR assay samples and detection of ubiquitylated and PARylated TOP3Bccs

DNA and RNA RADAR assay sample containing nucleic acids and covalent protein-nucleic acid adducts were recovered from HA or FLAG-tagged R338W TOP3B transfected or NSC690634 treated HCT116 cells as described previously(12,16,19). Aliquots of RADAR assay samples were saved as INPUT. The rest of the RADAR samples were diluted in IP buffer (50 mM Tris-HCl pH 7.4, 150 mM NaCl, 1 mM EDTA, 1% NP-40, 0.2% Triton X-100, 5% glycerol, 1 mM DTT, 20 mM N-ethylmaleimide and protease inhibitor cocktail) containing monoclonal Anti-TOP3B antibody [EP7779]—C-terminal (Abcam, CAT#: ab183520) or mouse monoclonal anti-FLAG M2 antibody (Millipore Sigma, St. Louis, MO, CAT#: F1804) or rabbit Anti-HA-Tag Monoclonal Antibody, Unconjugated, Clone C29F4 (Cell Signaling Technology, CAT#: 3724) and rotated overnight at 4°C. Next day, Pierce ChIP-grade Protein A/G Magnetic Beads was added and incubated with the samples for another 4 hr. Immunoprecipitated samples were washed with IP buffer twice, resuspended in TE buffer containing 5 mM CaCl2 and excess amount of micrococcal nuclease (Thermo Fisher Scientific, 100 units/μl). Samples were incubated in Thermomixer (1200 rpm, 37°C for 4 hr), resuspended in tris-glycine SDS sample buffer (Novex, LC2676) for SDS-PAGE and immunoblotted with different antibodies as indicated for detection of total TOP3Bccs and ubiquitylated or PARylated TOP3Bccs. In addition, each INPUT RADAR sample was subjected to slot-blotting and subjected to methylene blue staining to confirm equal DNA loading.

### 13. *In vitro* TOP3B PARylation assay

Recombinant TOP3B protein (10 ug) was incubated with recombinant PARP1 protein (10 ug), NAD+ (20-500 uM), 1 x PARylation buffer (50 mM Tris-HCl pH 8.0, 50 mM NaCl, 10 mM MgCl2, 2% glycerol, 1 mM DTT) and activated DNA. Following 20-min incubation at room temperature, the samples were subjected to Western blotting using anti-PAR and anti-TOP3B antibodies (44).

### 14. Identification of TOP3B ADP-ribosylation sites by LC-MS/MS

Recombinant TOP3B was PARylated *in vitro* as described in the previous section. *In vitro* PARylation assay samples (TOP3B +/-NAD+) were processed with trypsin/lysC digestion or chymotrypsin digestion and then purified by C18 desalting column. The cleaned-up peptides from each sample were dissolved in 15 μL of 0.1% formic acid, and 5 μL of each sample was analyzed in one 110-min LC-MS/MS run. The LC/MS/MS analysis of tryptic peptides for each sample was performed sequentially using a Thermo Scientific Orbitrap Exploris 480 Mass Spectrometer and a Thermo Vanquish Neo RSLCnano System. Peptides from trypsin digestion were loaded onto a peptide trap cartridge at a flow rate of 5 μL/min. The trapped peptides were eluted onto a reversed-phase Easy-Spray Column PepMap RSLC, C18, 2 μM, 100A, 75 μm × 250 mm (Thermo Scientific) using a linear gradient of acetonitrile (3-36%) in 0.1% formic acid. The elution duration was 60 min at a flow rate of 0.3 μl/min. Eluted peptides from the Easy-Spray column were ionized and sprayed into the mass spectrometer, using a Nano Easy-Spray Ion Source (Thermo) under the following settings: spray voltage, 1.6 kV, Capillary temperature, 275°C. Other settings were empirically determined. Raw data files were *search*ed against TOP3B1 protein sequence database using the Proteome Discoverer 3.1 software (Thermo, San Jose, CA) based on the SEQUEST algorithm. ADP-ribosylation (+541.061 Da) on N/D/R/K/E/C/T, Carbamidomethylation (+57.021 Da) of Cysteine, Oxidation (+15.995 Da) on Methionine M, and deamidation (+0.984 Da) on N, Q were set as dynamic modifications. The minimum peptide length was specified to be five amino acids. The precursor mass tolerance was set to 15 ppm, whereas fragment mass tolerance was set to 0.01 Da. The maximum false peptide discovery rate was specified as 0.01.

### 15. Site Directed Mutagenesis

Site-directed mutagenesis (SDM) was performed using Q5® Site-Directed Mutagenesis Kit (NEB, Cat# E0554S) following the manufacturer’s protocol, FLAG-tagged R338W TOP3B plasmid construct and following primer pairs:

T239A TOP3B,

FP:CAAACCAGAGGCGTACTGGGTGC,

RP:AAGGACTGGATTTTATCATGTCTC;

K246A TOP3B

FP:GCTGCAGGCCGCGGTTAACACTG,

RP:ACCCAGTAGGTCTCTG;

E282A TOP3B

FP:GCTGGAGAAGGCGGCCCAGGTGG,

RP:TTTGTCATGTTTAAAAACATCTGTGC;

E292A TOP3B

FP:AAGCAGGAAAGCGAAGGCCAAGC,

RP:GTGGCCTCCACCTGG;

K293A TOP3B

FP:CAGGAAAGAAGCGGCCAAGCAGAG,

RP:CTTGTGGCCTCCACC;

TOP3B K295A

FP:AGAAAAGGCCGCGCAGAGGCCCC,

RP:TTCCTGCTTGTGGCC;

E438A TOP3B

FP:AATTGGGCCCGCGCTCTTCACCT,

RP:CTGAAGGAGATGGTGCTCTGC

T488A TOP3B

FP:GGAGAAGCAGGCGAACCCACCCG,

RP:AGCATCTTCACCTCGCCCAC

N489A TOP3B

FP:GAAGCAGACGGCGCCACCCGACTAC,

RP:TCCAGCATCTTCACCTC

### 16. IP-mass spectrometry (IP-MS/MS) for detection of TOP3B interacting proteins

Co-IP-MS was carried out as previously described (16,18). Briefly, 10^7 cells were washed with PBS and incubated on a shaker with IP lysis buffer (50 mM Tris-HCl, pH = 7.4, 150 mM NaCl, 1 mM EDTA, 1 % NP-40, 0.2 % Triton X-100, 5 % glycerol, 1 mM DTT, 20 mM N-ethylmaleimide and protease inhibitor cocktail) to a final volume of 900 µL (supplemented with 2 µL benzonase (Sigma-Aldrich #E8263) and incubated on ice for 30 min, followed by sonication (thrice, 40 % power for 30 s pulse and 1 min rest) and clarified by centrifugation for 30 min at 4°C at 15k RCF. A fraction of the clarified supernatant was saved as input control, and the rest was diluted 3 to 4-fold in IP lysis buffer, with addition of anti-TOP3B (Abcam #AB_183520) or IgG antibodie**s** (3 µg/mL) and incubated overnight at 4°C with constant rotation. Next day, 1/30 volume of A/G magnetic beads (Thermo Fisher Scientific #26162) were added to the samples for additional incubation of 4 h. The magnetic beads were then collected and washed with RIPA buffer twice (150 mM NaCl, 1 % NP-40, 0.5 % sodium deoxycholate, 0.1 % SDS, 50 mM Tris, pH = 7.5, 1 mM DTT and protease inhibitor cocktail).

For MS analysis, the proteins were eluted from beads and digested with trypsin overnight using S-traps (Protifi) following the manufacturer’s protocol. Dried peptides were solubilized in 5 % acetonitrile and 0.05 % trifluoroacetic acid solution and applied to a trapping column and separated on a 75 μm × 15 cm, 2 μm Acclaim PepMap reverse phase column (Thermo Fisher Scientific) using an UltiMate 3000 RSLCnano HPLC (Thermo Fisher Scientific). Peptides were separated at a flow rate of 300 nL/min followed by online analysis by tandem mass spectrometry using a Thermo Orbitrap Exploris 480 mass spectrometer. Peptides were eluted into the mass spectrometer using a linear gradient from 4-35 % B over 120 min. Parent full-scan mass spectra were acquired at 120,000 FWHM resolution and product ion spectra at 15,000 resolution. Proteome Discoverer 3.0 (Thermo Fisher Scientific) was used to search the data against human proteins from the UniProt database using SequestHT. The search was limited to tryptic peptides, with maximally two missed cleavages allowed. Cysteine carbamidomethylation was set as a fixed modification, and methionine oxidation was set as a variable modification. The precursor mass tolerance was 10 ppm, and the fragment mass tolerance was 0.02 Da. The Percolator node was used to score and rank peptide matches using a 1% false discovery rate.

### 17. Excision of TOP3Bccs by recombinant FEN1

RADAR assay samples were prepared from HCT116 cells transfected with FLAG-tagged R338W TOP3B. Equal amount of RADAR samples were incubated with 500 nM recombinant human FEN1 in the presence or absence of FEN1 inhibitor or benzonase (250 units), or Micrococcal Nuclease (300 units) at 37°C for 30 min. Released TOP3B was detected by immunoblotting with anti-FLAG antibody after SDS-PAGE. Loading was tested by slot blotting and probing with methylene blue staining.

### 18. Quantification and Statistical Analysis

Quantifications were carried out using the Fiji software. Data are provided as means ± standard deviations (SD) from the number of independent experiments performed. Statistical analyses and graphical representation were carried out using GraphPad Prism 7 software. Statistical test methods are described in each figure legend. Statistical significance is indicated as follows: **P < 0.01, ***P < 0.001, ****P < 0.0001; n.s., not significant.

## AUTHOR CONTRIBUTIONS

S.S., Y.P., and Y.S. conceived the study. S.S. performed most of the biochemical and molecular biology experiments, with assistance from T.M. and L.Z. N.C.L. and G.M.W. performed the drug screening and analyzed the screening data. V.C., Y.Su., and J.C. performed the imaging experiments and analyzed the data. T.K.M. and L.M.J. performed mass spectrometry experiments and analyzed the data. B.W. and W.Z. contributed to biochemical studies. S.S. and Y.S. analyzed and interpreted the experimental results. Y.P. and Y.S. supervised the study and acquired funding. S.S. wrote the original manuscript. Y.P. and Y.S. revised the manuscript.

